# PICRUSt2-SC: an update to the reference database used for functional prediction within PICRUSt2

**DOI:** 10.1101/2025.01.27.635123

**Authors:** Robyn J. Wright, Morgan. G.I. Langille

## Abstract

**Summary:** PICRUSt2 is a bioinformatic tool that predicts microbial functions in amplicon sequencing data using a database of annotated reference genomes. We have constructed an updated database for PICRUSt2 that has substantially increased the number of bacterial (19,493 to 26,868) and archaeal (406 to 1,002) genomes as well as the number of functional annotations present. The previous PICRUSt2 database relied on many timely and computationally intensive manual processes that made it difficult to update. We constructed a new streamlined process to allow regular upgrades to the PICRUSt2 database on an ongoing basis, and used this process to create a new database, PICRUSt2-SC (Sugar-Coated). Additionally, we have shown that this updated database contains genomes that more closely match study sequences from a range of different environments. The genomes contained in the database therefore better represent these environments and this leads to an improvement in the predicted functional annotations obtained from PICRUSt2.

**Availability:** PICRUSt2 source code is freely available at https://github.com/picrust/picrust2 and at https://anaconda.org/bioconda/picrust2. The latest version of PICRUSt2 at the time of writing is also archived: https://doi.org/10.5281/zenodo.15119781. The PICRUSt2-SC database comes pre-installed with PICRUSt2 from version 2.6.0 onwards. Step-by-step instructions for making the updated database are at https://github.com/picrust/picrust2/wiki/Updating-the-PICRUSt2-database. All code used for the analyses and figures in this manuscript is at https://github.com/R-Wright-1/PICRUSt2-SC_application_note and https://doi.org/10.5281/zenodo.15119770.

## 1. Introduction

PICRUSt2 (**P**hylogenetic **I**nvestigation of **C**ommunities by **R**econstruction of **U**nobserved **St**ates) is a widely used bioinformatic tool that allows the prediction of microbial community functions based on amplicon sequencing data. The first version of PICRUSt (PICRUSt1) was published in 2013 [1] and the second version (PICRUSt2) was published in 2020 [2]. In PICRUSt2, 16S rRNA gene sequences collapsed to amplicon sequence variants (ASVs) are placed into a reference tree using multiple-sequence alignment and then hidden-state prediction is used to infer the gene content of study sequences based on the gene content of the taxa used to construct the reference tree. Information on their predicted 16S rRNA gene copy number and their abundance in the original samples is used to weight the predicted genome annotations to generate a predicted metagenome.

One of the largest limitations to PICRUSt2 currently is that the default database uses functional annotations acquired from the Integrated Microbial Genomes database [3] in 2017. The number of genomes and functions have continued to expand during this time and timely updates to PICRUSt2 are essential for its accuracy and functional comprehensiveness. However, the previous approach for updating the default database and functions was difficult due to a reference tree needing to be re-built with newly added genomes. We have constructed a new database for PICRUSt2, PICRUSt2-SC (**S**ugar-**C**oated), that uses Genome Taxonomy Database (GTDB) r214 genomes [4–7] and the bacterial and archaeal phylogenetic trees released with GTDB. We have annotated these genomes using Eggnog [8, 9], and as the GTDB genomes are readily available for download, we have provided instructions for PICRUSt2 users to add annotations to this updated database.

## 2. PICRUSt2-SC Database creation

All genomes (*n*=402,709) from the GTDB r214.1 were downloaded from the GTDB repository. Only representative genomes that were present in the GTDB bacterial or archaeal phylogenetic trees, had a high quality 16S rRNA gene and that were ≤ 10% contamination and ≥ 90% completion were used, giving *n*=27,870 total genomes (*n*=26,868 and *n*=1,002 genomes for bacteria and archaea, respectively). Full details of the methods used for 16S rRNA gene identification and processing are in Supplementary Information section 2.1. This has increased the number of taxa present at every phylogenetic rank from phylum to species (Fig. 1), with approximately 3-fold or 4-fold more species present in the PICRUSt2-SC database for bacteria or archaea, respectively, compared with the previous PICRUSt2 database (hereafter referred to as PICRUSt2-oldIMG). Included bacterial genomes have a median completeness of 99.35% and contamination of 0.66% while archaeal genomes have a median completeness of 99.03% and contamination of 0.67%.

**Figure 1.**
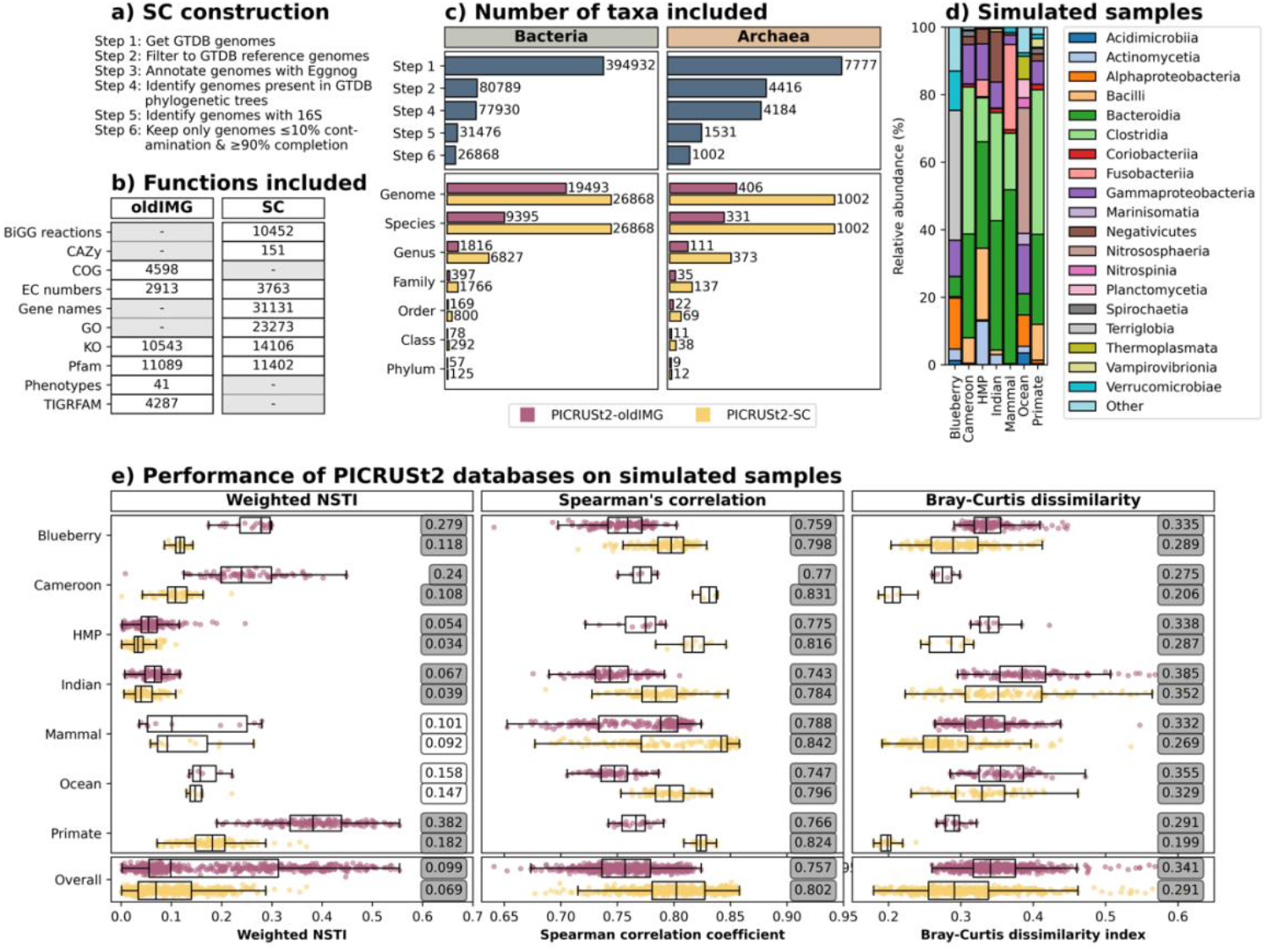
Comparison of the PICRUSt2-oldIMG and PICRUSt2-SC databases showing: (**a**) the steps in the construction of the PICRUSt2-SC database; (**b**) the number of functions annotated within different frameworks for the default and updated databases (note that not all frameworks were included in both databases); (**c**) the number of taxa included for each step of the PICRUSt2-SC construction (top) and for each phylogenetic rank (bottom) for bacteria and archaea; (**d**) composition at the class level for the simulated samples (the mean relative abundance is shown for each dataset); and (**e**) the performance of the PICRUSt2-oldIMG and PICRUSt2-SC databases on the simulated samples from each dataset and overall (bottom). Spearman’s correlation and Bray-Curtis dissimilarity is shown for KEGG orthologs (EC numbers are in Fig. S5). Individual points are shown for each sample with points being coloured pink for the PICRUSt2-oldIMG database and yellow for the PICRUSt2-SC database. Boxplots represent the median, upper and lower quartiles and whiskers show the range of the data (1.5 times the Interquartile Range) and values in boxes are medians. The results for t-tests between the PICRUSt2-oldIMG and PICRUSt2-SC are shown with grey shading for significant (p ≤ 0.05) tests.

Annotation of all representative genomes within the new PICRUSt2-SC database was carried out using Eggnog v2.1.12 with Prodigal v2.6.3 [10] for gene prediction and the Eggnog diamond database v5.0.2. This gave Biochemical, Genetic and Genomic (BiGG) model reactions [11], Carbohydrate Active Enzymes (CAZy) [12], Enzyme Commission (EC) number, Gene Ontology (GO) [13, 14], Kyoto Encyclopedia of Genes and Genomes (KEGG) orthologs (KO) [15], Protein family (Pfam) [16], and gene name annotations for all genomes. The default PICRUSt2 database contained annotations for KO, EC, Pfam, Clusters of Orthologous Genes (COG) [17, 18], Phenotype [3] and TIGRFAM [19] annotations. By default, only the KO and EC annotations are used, as in the previous version. In the PICRUSt2-SC database, we have 1.3-fold more annotations than the PICRUSt2-oldIMG database for both KOs (14,106 *versus* 10,543) and EC numbers (3,763 *versus* 2,913), while the number of Pfam annotations remains similar (Fig. 1). See supplementary information section 2.2 for further details. Users may also add their own functions to the new database and we provide instructions for doing so on the PICRUSt2 Github.

## 3. Placement of sequences into phylogenetic trees

With the PICRUSt2-oldIMG database, study sequences are placed in a reference phylogenetic tree constructed using 16S rRNA gene sequences that contains both bacterial and archaeal sequences. In our PICRUSt2-SC database, we now use separate (unrooted) trees for bacteria and archaea constructed using multiple marker genes (from GTDB), and we therefore initially place sequences in both trees and perform 16S copy number prediction and Nearest Sequenced Taxon Index (NSTI) calculations for both trees. We then compare the NSTI obtained for each and choose the most appropriate (lowest NSTI) for functional predictions. These updates to the code used for running PICRUSt2 are described in Supplementary Information section 3.1. To verify that this tree placement is appropriate, we used the same 16S datasets used in Douglas *et al*. [2]: (i) Blueberry *n*=22, bulk soil and blueberry rhizosphere samples [20, 21]; (ii) Cameroon, *n*=57, stool samples from Cameroonian individuals [22, 23]; (iii) HMP, *n*=137, samples spanning the human body from the Human Microbiome Project [24]; (iv) Indian, *n*=91, stool samples from Indian individuals [25]; (v) Mammal, *n*=8, mammalian stool samples [26]; (vi) Primate, *n*=77, non-human primate stool samples [27]; and (vii) Ocean, *n*=6, ocean samples [28]. We took the processed feature tables and representative sequences used by Douglas *et al*. and classified these taxonomically using the naïve bayes Scikit-Learn classifier trained on full-length 16S rRNA gene sequences from the Greengenes2 database [29] within QIIME2 (v2024.5) [30]. We ran PICRUSt2 using both the PICRUSt2-oldIMG and the PICRUSt2-SC databases for each of the datasets.

For PICRUSt2-SC predictions, the domain obtained from the taxonomic classifications and the domain of the tree giving the lowest NSTI value for each sequence agreed for 21,543 of 21,552 (99.9%) total sequences within these datasets. See Supplementary Information section 3.2 for details on the nine ASVs that were potentially placed into the wrong tree. Comparing the NSTIs obtained with the PICRUSt2-SC *versus* the PICRUSt2-oldIMG database on a per sequence basis and on a per sample basis (*i*.*e*., the weighted NSTIs that account for sequence abundance within samples), we find that the NSTIs have decreased significantly (t- test *p* ≤ 0.05) in almost all cases (Supplementary Information section 3.3).

## 4. Verification of the functional predictions obtained with the updated database

The functional predictions obtained with the PICRUSt2-SC database were correlated with those obtained with the PICRUSt2-oldIMG database (Supplementary Information section 4.1). In Douglas *et al*. [2], Spearman correlation coefficients were calculated between PICRUSt2-obtained KO predictions and HUMAnN2-obtained KO annotations of the paired metagenome samples. Although we did not expect this to be a reliable standard for a database constructed at a different time using different databases for annotation, we calculated Spearman correlation coefficients and Bray-Curtis dissimilarity indices between the PICRUSt2-predictions for both databases and the HUMAnN2 annotations for comparability with the previous study (Supplementary Information section 4.2).

We instead constructed mock samples using Eggnog-annotated genomes with simulated abundances. Briefly, we matched the taxa in the seven datasets described above to GTDB genomes that were not present in the PICRUSt2-SC database and used these to construct mock communities that matched the abundances within the seven real datasets. Full details of this mock sample construction can be found in Supplementary Information section 4.3. The mock samples that we constructed comprised a range of taxa typical of the environment that they were designed to simulate (Fig. 1), for example, the Blueberry soil samples contained genomes from the Terriglobia class, while the Ocean water samples contained genomes from the Nitrososphaeria class. We ran PICRUSt2 with both the PICRUSt2-oldIMG and the PICRUSt2-SC databases using the mock representative sequences and feature tables for each dataset. Comparing the NSTIs obtained with the PICRUSt2-SC database *versus* the PICRUSt2-oldIMG database on a per sample basis (*i*.*e*., the weighted NSTIs that account for sequence abundance within samples), we find that the NSTIs have decreased in all cases, most of which are statistically significant (Fig. 1). The NSTIs have decreased from a median of 0.099 (range 0.054 [HMP] – 0.382 [Primate]) with PICRUSt2-oldIMG to 0.069 (range 0.034 [HMP] – 0.182 [Primate]) with PICRUSt2-SC. While the median NSTI per sequence (Supplementary Information section 4.4) was slightly lower with the PICRUSt2-oldIMG database for the Cameroon and Indian datasets, the median weighted NSTIs were always lower with the PICRUSt2-SC database, suggesting that it is rarer sequences that have higher NSTIs.

We compared the functional predictions obtained using PICRUSt2 with either the PICRUSt2-oldIMG or the PICRUSt2-SC database with the gold-standard and calculated both Spearman correlation coefficients and Bray-Curtis dissimilarity indices. Median Spearman correlation coefficients are significantly higher for the PICRUSt2-SC (median 0.802; range 0.784 [Indian] – 0.842 [Mammal]) *versus* the PICRUSt2-oldIMG (median 0.757; range 0.743 [Indian] – 0.788 [Mammal]) database for both KOs (Fig. 1) and EC numbers (Supplementary Information section 4.4).

Bray-Curtis dissimilarity indices were also significantly lower for the PICRUSt2-SC (median 0.291; 0.352 [Indian] – range 0.199 [Primate]) *versus* the PICRUSt2-oldIMG (median 0.341; 0.385 [Indian] – range 0.275 [Cameroon]) database for both KOs (Fig. 1) and EC numbers (Supplementary Information section 4.4). KOs had both higher Spearman correlation coefficients and lower Bray-Curtis dissimilarity indices between the mock and PICRUSt2-predictions than EC numbers, and the lowest correlations and highest Bray-Curtis dissimilarity indices were found consistently with the Indian dataset.

## 5. Computational resources required to run the updated database

We used the sequences within the 16S datasets described above to simulate datasets with 10, 100, 1,000 or 10,000 sequences and 10, 100 or 1,000 samples (Supplementary Information section 5). Because of the increase in the number of genomes and annotations, the updated PICRUSt2-SC database requires more time and memory to run than PICRUSt2-oldIMG; a dataset containing 1,000 sequences and 1,000 samples being run using 12 threads now takes ∼23 mins to run and used 36 GB RAM (compared with ∼12 mins and 27 GB RAM with PICRUSt2-oldIMG; Table S1). These memory requirements can be further reduced with modifications to some of the parameters used to run and we believe that the resources required are still reasonable for most researchers.

## 6. Conclusions

We have shown that the updated PICRUSt2-SC database using GTDB genomes performs as expected and outperforms the previous PICRUSt2-oldIMG database in almost all cases, although PICRUSt2-SC requires more time and computational resources than PICRUSt2-oldIMG. This provides PICRUSt2 users with a database that contains more genomes and functions and provides a framework for the PICRUSt2 database to be updated more frequently in the future. The relative ease with which users may now download the database genomes allows users to add their own functions of interest to the database and opens the door for new and improved methods for functional annotations to be used, such as those being developed that use Artificial Intelligence. It remains important for PICRUSt2 users to remember that the output of PICRUSt2 contains functional predictions that are based on the functional annotations of the genomes that are present in the database, but large differences in the genomic content of even very closely related strains with identical 16S rRNA gene sequences may be possible. PICRUSt2-generated functional predictions are therefore recommended as a hypothesis-generating tool and not as the sole piece of evidence for associations between a function and biological phenomena.

## Funding and acknowledgements

This work was supported by an NSERC Discovery Grant (2016-05039) and was enabled in part by the Digital Research Alliance of Canada (alliancecan.ca).

We thank Dr. Gavin Douglas for invaluable insight and assistance with troubleshooting and several PICRUSt2 users for their time testing and giving feedback on an early version of the PICRUSt2-SC database.

