## Supplementary Information and Figures for "PICRUSt2-SC: an update to the reference database used for functional prediction within PICRUSt2"

### PICRUSt2-SC supplementary information

#### 2. Database creation and pipeline updates

##### 2.1 Genome curation and 16S rRNA gene identification

For genomes to be included in the new PICRUSt2-SC database, we required them to be present in the GTDB bacterial or archaeal phylogenetic trees (*n*=77,930 or *n*=4,184, respectively) and to have at least one 16S rRNA gene (hereafter referred to as 16S) copy. To identify 16S genes within genomes, Barrnap v0.9 [1] was run with the bacterial or archaeal models on the bacterial or archaeal genomes, respectively. Sequences shorter than 80% of the full length (~1500 bp) 16S gene were rejected. This left *n*=32,595 or *n*=1,553 bacterial or archaeal genomes, respectively. As in Douglas *et al.* [2], where multiple 16S copies were present in a single genome, the centroid was identified using the VSEARCH (v2.4.4) [3] cluster_fast command with 90% identity. Where multiple centroids were found, a single one was chosen at random (using the Python package random). To identify genomes with identical 16S genes, the VSEARCH cluster_fast command was again used, but this time with 100% identity. Alignment of all 16S genes was carried out using ssu-align [4, 5] separately for bacteria and archaea, and 16S genes that did not match the model for the domain that they were supposed to come from were discarded. This gave *n*=31,476 or *n*=1,531 genomes with 16S genes for bacteria or archaea, respectively, for use in the PICRUSt2-SC database.

For genomes that had identical 16S rRNA genes, the genome with the highest completion percentage (estimated as part of the GTDB database construction using CheckM [6]) was chosen for trait prediction. Only genomes with ≤ 10% contamination and ≥ 90% completion were used, giving *n*=27,870 total genomes (*n*=26,868 and *n*=1,002 genomes for bacteria and archaea, respectively). GTDB tree files were filtered to have only the included genomes using the Python packages ete3 [7] and Biopython [8] and the model files needed for tree placement were created using RAxML [9] and RAxML-NG [10].

##### 2.2 Annotation

As in the PICRUSt2-oldIMG database, copy numbers higher than 10 in this final set were re-coded to be 10 to decrease the number of possible prediction states. Like for the PICRUSt2-oldIMG database, we have taken the files for mapping EC numbers to MetaCyc pathways and for MetaCyc reactions from the latest HUMAnN version (HUMAnN3) [11]. Mapping from KOs to pathways or modules remains possible with the existing mapping files; these are taken from the last publicly available mapping, and users may update this file themselves if they have a KEGG database subscription.

#### 3. Placement of sequences into phylogenetic trees

##### 3.1 PICRUSt2 pipeline updates

Previously, the default use of PICRUSt2 involved using the picrust2_pipeline.py script to run the key steps within PICRUSt2: (i) placement of study sequences into the reference tree; (ii) hidden-state prediction for 16S copy number and NSTI calculations as well as gene family abundances; (iii) metagenome prediction; and (iv) inference of pathway abundances. We have not made major updates to this but in the picrust2_pipeline.py script in PICRUSt2 v2.6.0 the steps (i) and (ii) will now be run separately for bacteria and archaea and the predicted functional abundances will be combined for metagenome prediction and the inference of pathway abundances. This means that step (i) is now split to: (a) placement of study sequences into the bacterial reference tree; and (b) placement of study sequences into the archaeal reference tree. Step (ii) is split to: (a) hidden-state prediction for 16S copy number and NSTI calculations using the bacterial reference tree; (b) hidden-state prediction for 16S copy number and NSTI calculations using the archaeal reference tree; (c) comparison of NSTI values in the bacterial and archaeal reference trees for all study sequences. The domain with the lowest NSTI value is chosen and the files for each domain are filtered to only contain study sequences for which that domain is the best fit; (d) hidden-state prediction for gene family abundances, separately for bacteria and archaea; and (e) bacterial and archaeal predictions are combined for each trait/gene family into a single file.

We have also added a column to the output file that details which reference genome was closest to each study sequence in the phylogenetic tree, allowing users to verify the closest taxonomy to each study sequence. These updates have not changed any of the methods used by PICRUSt2, and the previous default use of PICRUSt2 remains in the source code (picrust2_pipeline_oldIMG.py) so that users may choose to use the PICRUSt2-oldIMG database for comparability with previous studies.

##### 3.2 Comparison of domain obtained by QIIME2 taxonomic classifications and tree used by PICRUSt2 for metagenome inference

The remaining ASVs came from either the Ocean (*n*=8) or Primate (*n*=1) datasets (both Illumina MiSeq datasets). Five of the Ocean ASVs had NSTI >1 in both trees, while three were classified as belonging to the Woesearchaeales order (UBA12501, UBA583 and B72-G16 families) and had NSTI 0.13-0.21 in the bacterial tree and 1.66-1.89 in the archaeal tree. While genomes belonging to the Woesearchaeales order are present in the archaeal tree, genomes from the families that these ASVs were classified as were not included due to low genome completeness. Furthermore, NCBI BLAST [12] searches for these sequences revealed low sequence similarity with reference databases (90-95% identity with both uncultured bacterial and uncultured archaeal taxa), suggesting that these sequences may be ambiguous. The

ASV from the Primate dataset had NSTIs of 0.11 and 0.19 in the bacterial and archaeal trees, respectively, but was classified taxonomically as *Cenarchaeum* (Nitrososphaeria_A, Archaea). Genomes from this genus are present in the archaeal tree, so it is not clear why this misclassification has occurred, however, the ASVs within the Primate dataset are much shorter than is typically used for modern amplicon sequencing studies (150 bp). These misclassifications occur at a very low rate, but as they are all from ASVs classified as archaea, we recommend that if datasets are made up of a large proportion of archaea, users further verify that the insertions of sequences into the tree within PICRUSt2-SC seem appropriate.

##### 3.3 Comparison of NSTIs for the PICRUSt2-oldIMG *versus* PICRUSt2-SC databases

The per sequence NSTIs have decreased on average 0.186 (median 0.075) while the per sample NSTIs have decreased on average 0.091 (median 0.053) across all datasets (Fig. S1A). The median NSTIs with the PICRUSt2-oldIMG database ranged between 0.05 (HMP) and 0.32 (Primate) and with the PICRUSt2-SC database between 0.03 (HMP) and 0.15 (Ocean) (Fig. 2a). The medians have changed by -0.07 (Blueberry, t-test $p < 0.001$), -0.14 (Cameroon, $p < 0.001$), -0.02 (HMP, $p < 0.001$), +0.02 (Indian, $p < 0.001$), -0.01 (Mammal, $p < 0.001$), -0.13 (Ocean, $p < 0.001$) and -0.19 (Primate, $p < 0.001$), suggesting that the PICRUSt2-SC database better covers the diversity present in these real datasets. Comparing the NSTIs on a *per* sample basis (*i.e.*, the weighted NSTIs that account for ASV abundance within samples), we find larger differences between the PICRUSt2-oldIMG and the PICRUSt2-SC database (Fig. S1B). The median weighted NSTIs with the PICRUSt2-oldIMG database ranged between 0.05 (HMP) and 0.38 (Primate) and with the PICRUSt2-SC database between 0.03 (HMP) and 0.18 (Primate). The median weighted NSTIs have changed by -0.16 (Blueberry, t-test $p < 0.001$), -0.13 (Cameroon, $p < 0.001$), -0.02 (HMP, $p < 0.001$), -0.03 (Indian, $p < 0.001$), -0.01 (Mammal, N.S.), -0.01 (Ocean, N.S.) and -0.20 (Primate, $p < 0.001$).

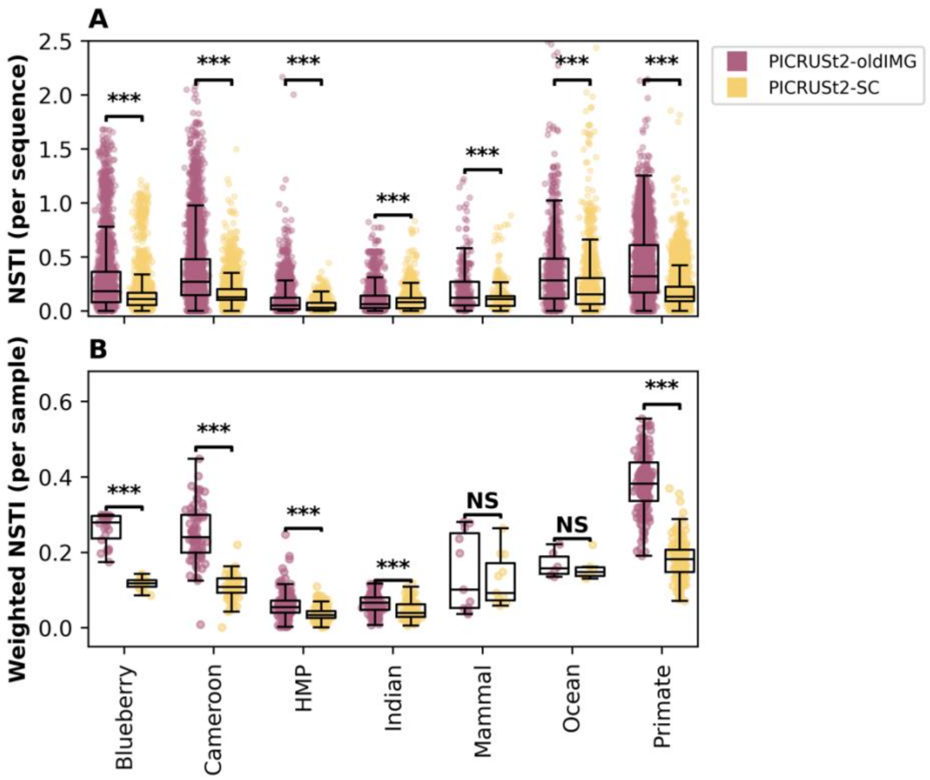


**Figure S1.** Nearest Sequenced Taxon Indices (NSTIs) for the seven datasets used in Douglas *et al.* for the PICRUSt2-oldIMG (pink) or PICRUSt2-SC (yellow) databases. NSTIs are shown with individual points per sequence in (**A**) and per weighted NSTIs within a sample in (**B**). Boxplots represent the median, upper and lower quartiles and whiskers show the range of the data (1.5 times the Interquartile Range). The results for t-tests between the PICRUSt2-oldIMG and PICRUSt2-SC results are shown, with *** denoting $p < 0.001$ and NS denoting $p > 0.05$.

#### 4. Verification of the functional predictions obtained with the PICRUSt2-SC database

##### 4.1 Correlation between PICRUSt2-oldIMG and PICRUSt2-SC database predictions

The Spearman correlation coefficients between functional predictions obtained with both databases were statistically significant ($p \leq 0.001$) and high (median 0.89 [Indian] – 0.93 [Ocean], 0.85 [Indian] – 0.90 [Blueberry] and 0.85 [Mammal] – 0.90 [Cameroon] for EC numbers, KOs and Metacyc pathways, respectively; Fig. S2).

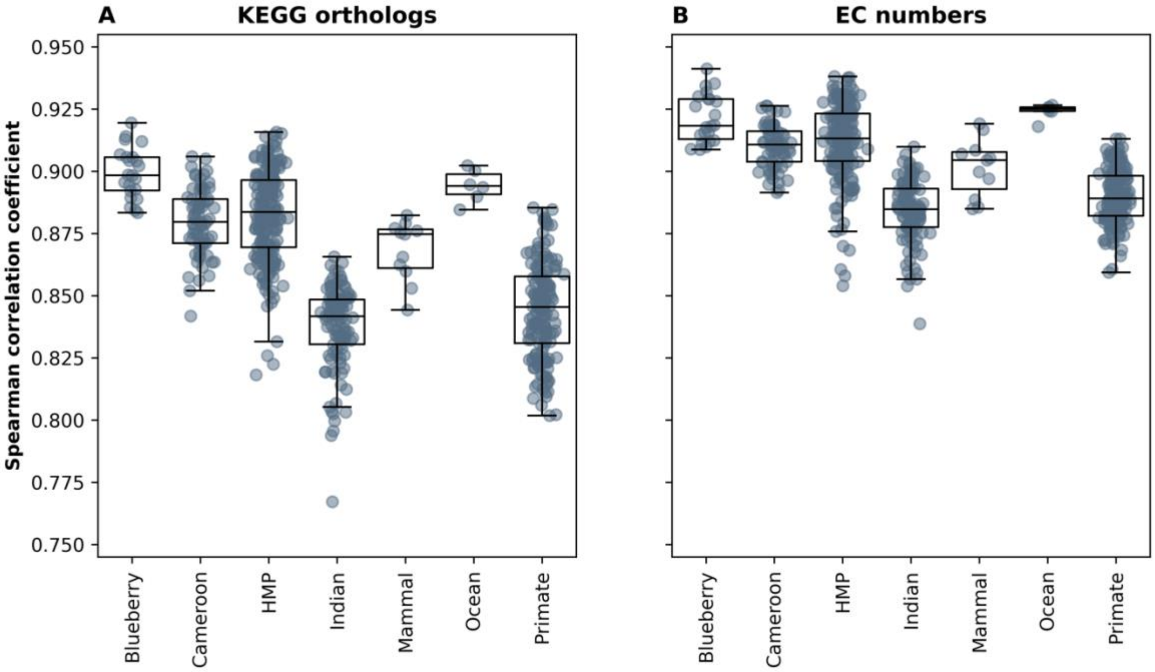


**Figure S2.** Spearman correlation coefficients between functional predictions obtained using the PICRUSt2-oldIMG database and the PICRUSt2-SC database for (**A**) EC numbers and (**B**) KOs. Only EC numbers or KOs that appeared in both sets of functional predictions were included in these calculations. Each sample is represented as a point with boxplots representing the median, upper and lower quartiles. Whiskers show the range of the data (1.5 times the Interquartile Range).

##### 4.2 Comparison of the PICRUSt2-SC database with the "gold-standard" used by Douglas *et al.*

In Douglas *et al.* [2], PICRUSt2 with the PICRUSt2-oldIMG database was validated by comparing Spearman's correlation coefficients between PICRUSt2-obtained KEGG ortholog (KO) predictions with a "gold-standard" that used HUMAnN2-obtained KO annotations of the MGS samples. We calculated Spearman correlation coefficients as well as Bray-Curtis dissimilarity indices between the PICRUSt2-predictions for both databases and the MGS HUMAnN2 annotations. Median Spearman correlation coefficients for KOs and EC numbers were slightly lower for the PICRUSt2-SC database than for the PICRUSt2-oldIMG database for all datasets aside from the Primate dataset (Fig. S3AB). Median Bray-Curtis dissimilarity indices were very similar for both databases, with the PICRUSt2-SC database performing slightly better on the EC numbers and the PICRUSt2-oldIMG database on KOs (Fig. S3CD).

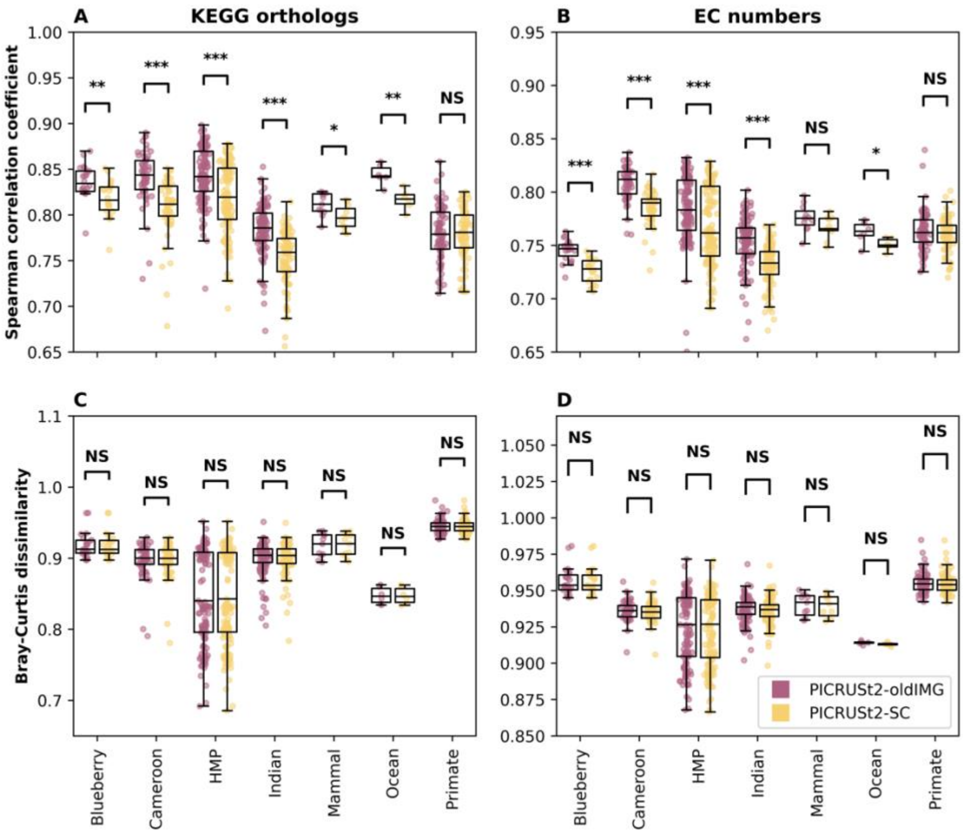


**Figure S3.** Spearman correlation coefficients (A, B) and Bray-Curtis distances (C, D) between "gold-standard" shotgun metagenome functional annotations and PICRUSt2 predictions for EC numbers (A, C) and KOs (B, D) for the PICRUSt2-oldIMG (pink) and PICRUSt2-SC (yellow) databases. Each sample is represented as a point with boxplots representing the median, upper and lower quartiles. Whiskers show the range of the data (1.5 times the Interquartile Range). The results for t-tests between the PICRUSt2-oldIMG and PICRUSt2-SC results are shown, with *** denoting $p < 0.001$, ** denoting $p < 0.01$, * denoting $p \leq 0.05$, and NS denoting $p > 0.05$.

#### 4.3 Construction of the simulated/mock samples

We collapsed the taxonomic classifications obtained above to the genus rank (where possible) and collapsed the feature tables at this rank. This gave 1,480 unique taxa, 1,179 of which had classifications at the genus rank. We then picked a genome from the GTDB database to match each of these taxa as best as possible; we were aiming to construct communities that contained a realistic number of species that represented diverse habitats, but not that were exactly as might be found in the environment. The genomes that we picked were: (i) not a representative species (*i.e.,* not present in the PICRUSt2-SC database); (ii) ≥90% complete; (iii) ≤10% contaminated; (iv) the same taxonomy as obtained from the samples to the lowest rank feasible.

We identified 16S copies in the genomes using the same methods described above, giving 811 genomes with at least one 16S copy. This gave 214, 211, 166, 176, 41, 254, and 266 taxa per dataset for the Blueberry, Cameroon, HMP, Indian, Mammal, Ocean, and Primate datasets, respectively. Where multiple 16S copies exist within genomes, these were randomly sampled to make up the observed sequence counts within samples. Each dataset was then processed in one of two ways: (i) 16S sequences were left untrimmed (mean 1,175, 1,268, 1,394, 1,336, 1,239, 1,302, 1,227 bp, for the datasets, respectively; range 396-1,590 bp); (ii) 16S sequences were trimmed to the V4-V5 region (Universal V4-V5 primers [13]) using cutadapt [14]. For both cases, sequences were clustered at 100% identity using VSEARCH to generate abundance tables of representative sequences.

##### 4.4 Performance of the databases on the simulated samples

The per ASV NSTIs have decreased significantly (t-test $p \leq 0.05$) from a median of 0.027 with the PICRUSt2-oldIMG database (range 0.02 [Cameroon] – 0.051 [Blueberry]) to 0.024 (range 0.021 [HMP/Mammal] – 0.029 [Ocean]) with the PICRUSt2-SC database (Fig. S4). The per sample NSTIs (weighted NSTIs) are discussed in the main text.

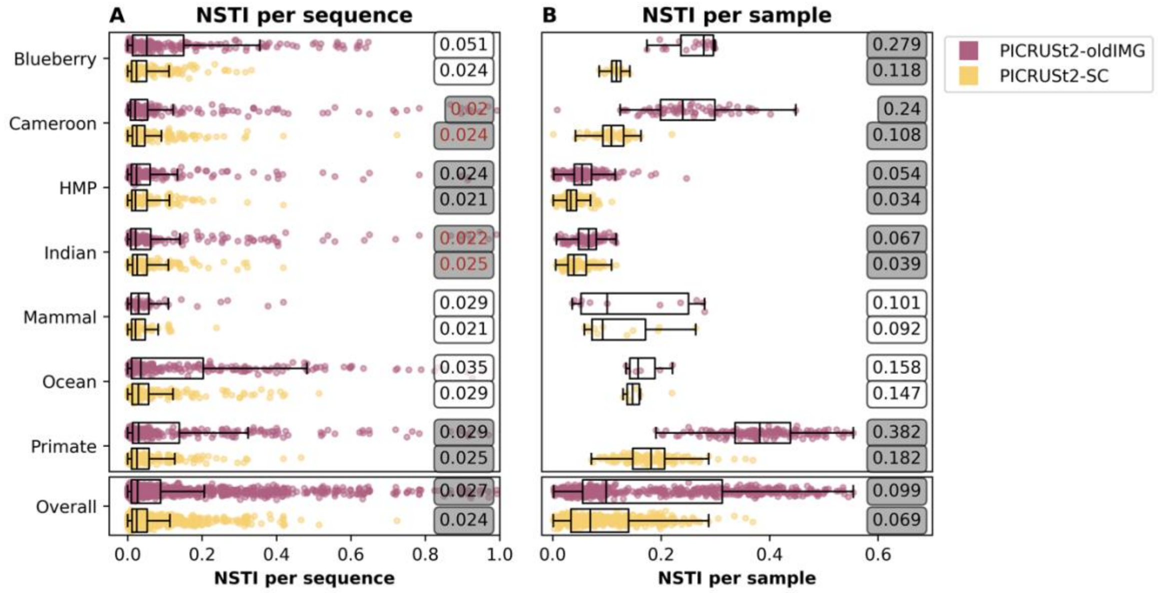


**Figure S4**. Nearest Sequenced Taxon Indices (NSTI) in the mock samples for the PICRUSt2-oldIMG (pink) and PICRUSt2-SC (yellow) databases. Each point shown is a sequence (left) or sample (right) within the dataset. Each point shown is a sample within a dataset. Boxplots represent the median, upper and lower quartiles and whiskers show the range of the data (1.5 times the Interquartile Range). Red text indicates that the PICRUSt2-oldIMG database outperformed the PICRUSt2-SC database. The results for t-tests between the PICRUSt2-oldIMG and PICRUSt2-SC are shown with grey shading for significant ($p \leq 0.05$) tests.

KOs had both higher Spearman correlation coefficients and lower Bray-Curtis distances between the mock and PICRUSt2-predictions than EC numbers (KOs are shown in Fig. 1 and EC numbers in Fig. S5). For the EC numbers, correlations ranged between 0.396 (Blueberry) – 0.502 (Mammal) for the PICRUSt2-oldIMG database and 0.453 (Primate) – 0.521 (Mammal) for the PICRUSt2-SC database while Bray-Curtis dissimilarity indices ranged between 0.612 (Indian) – 0.512 (Ocean) and 0.571 (Indian) – 0.44 (Ocean). Spearman's correlation coefficients and Bray-Curtis dissimilarity indices are shown for KOs in the main text. In all cases, the PICRUSt2-SC performed significantly (t-test $p \leq 0.05$) better than PICRUSt2-oldIMG.

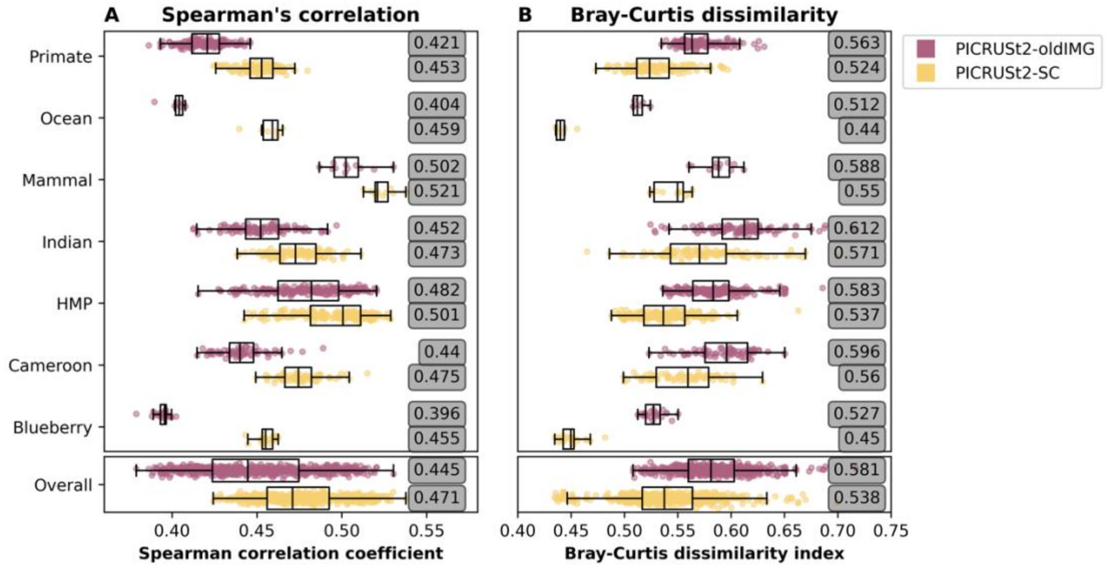


**Figure S5.** Spearman correlation coefficients and Bray-Curtis distances between mock community and PICRUSt2 predictions on full-length 16S sequences for EC numbers for the PICRUSt2-oldIMG (pink) and PICRUSt2-SC (yellow) databases. Each sample is represented as a point with boxplots representing the median, upper and lower quartiles. Whiskers show the range of the data (1.5 times the Interquartile Range). The results for t-tests between the PICRUSt2-oldIMG and PICRUSt2-SC are shown with grey shading for significant ($p \leq 0.05$) tests.

##### 4.5 Effect of trimming 16S sequences to the V4-V5 region

Trimming the 16S sequences within the mock community to only the V4-V5 region (described above) made very little difference to the Spearman correlation coefficients or Bray-Curtis distances between the predictions and the mock communities (Fig. S6).

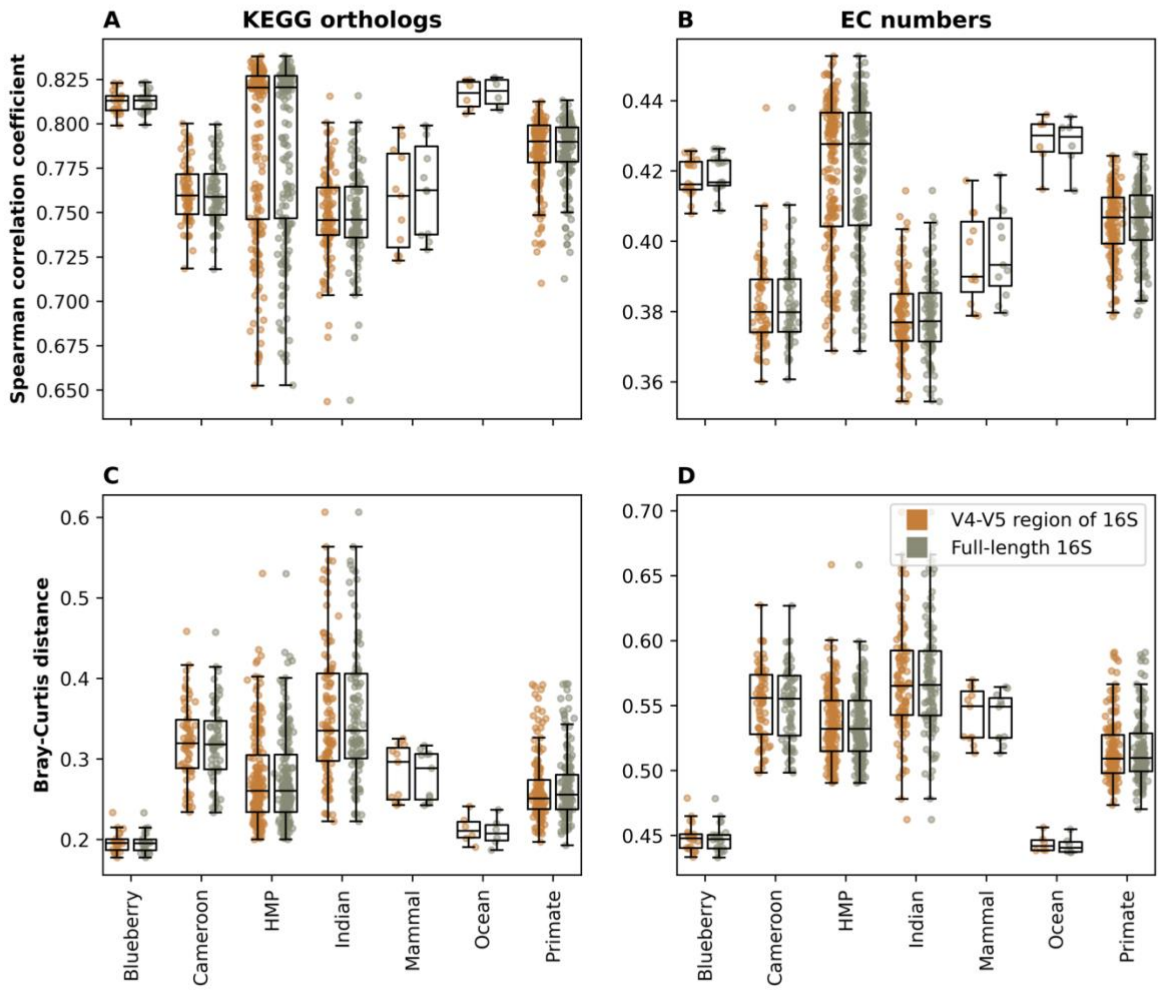


**Figure S6.** Spearman correlation coefficients (A, B) and Bray-Curtis distances (C, D) between mock community and PICRUSt2 predictions obtained using the PICRUSt2-SC database on V4-V5 region (orange) or full-length 16S (green) sequences for KOs (A, C) and EC numbers (B, D). Each sample is represented as a point with boxplots representing the median, upper and lower quartiles. Whiskers show the range of the data (1.5 times the Interquartile Range).

#### 5. Computational resources used to run PICRUSt2-oldIMG and PICRUSt2-SC

To determine the additional resources used to run the updated PICRUSt2-SC database compared with the previous PICRUSt2-oldIMG we constructed several simulated datasets. Sequence files with 10, 100, 1,000 or 10,000 sequences were constructed by taking sequences from the real datasets at random, and feature tables were constructed with 10, 100 or 1,000 samples for each of the sequence files by assigning the sequences with random abundances. These datasets were then run using both PICRUSt2-oldIMG and PICRUSt2-SC using the default parameters with 12 threads (unless stated otherwise) on a server running Ubuntu 20.04 with 1.5 TB RAM and 4 × Intel(R) Xeon(R) CPU E7-8870 v4 @ 2.10GHz=40 cores (80 threads).

For both PICRUSt2-oldIMG and PICRUSt2-SC, the number of samples included has little impact on the maximum memory (RAM) usage but does increase the amount of time required to run (Table S1). Increasing the number of sequences within the dataset increases the memory usage, with an increase from 10 to 10,000 sequences doubling the memory used for PICRUSt2-oldIMG and using ~1.7x more memory for PICRUSt2-SC. PICRUSt2-SC takes approximately double the time that PICRUSt2-oldIMG takes to run, and ~1.6x more memory for 10 sequences or ~1.3x more memory for 1,000 sequences. While this is a substantial increase, we do not believe that it will preclude most analyses from being carried out as these are still modest requirements compared with what is required for many steps in the bioinformatic analysis of microbiomes. Furthermore, while the figures we have presented (Table S1) were determined on a server, it is worth noting that these memory resources are not necessarily a requirement for running PICRUSt2, and we have successfully run PICRUSt2 (oldIMG and SC) on a 2020 MacBook Air with a M1 chip and 16 GB RAM with the default settings. Under these circumstances, PICRUSt2 will take longer to run but should still be able to run. Memory use can be further reduced within several steps of the pipeline by setting the chunk_size within the sequence placement and hidden-state prediction steps. The time and memory use given below are therefore used for comparison between the two databases and the effect of increasing sample or sequence numbers on these.

**Table S1.** Time and computational resources required to run PICRUSt2-oldIMG and PICRUSt2-SC.

| Database | Samples | PICRUSt2-oldIMG | | | | | PICRUSt2-SC | | | | |
|---|---|---|---|---|---|---|---|---|---|---|---|
| | | Sequences | | | | | Sequences | | | | |
| | | 10♥ | 10 | 100 | 1000 | 10000 | 10♥ | 10 | 100 | 1000 | 10000 |
| User time (min)* | 10 | 49 | 46 | 47 | 50 | 87 | 83 | 91 | 87 | 94 | 168 |
| | 100 | 48 | 49 | 48 | 52 | 91 | 85 | 89 | 93 | 95 | 175 |
| | 1000 | 65 | 63 | 66 | 73 | 124 | 115 | 117 | 126 | 139 | 247 |
| Elapsed time (min:sec)^ | 10 | 46:28 | 7:01 | 7:03 | 8:17 | 17:42 | 78:47 | 15:30 | 13:15 | 14:58 | 51:53 |
| | 100 | 44:48 | 7:38 | 7:16 | 8:28 | 19:08 | 81:04 | 13:45 | 16:24 | 15:29 | 55:11 |
| | 1000 | 62:41 | 9:08 | 9:46 | 12:07 | 39:26 | 112:05 | 15:51 | 17:34 | 22:36 | 98:12 |
| Memory usage (GB)♦ | 10 | 17.14 | 17.14 | 23.36 | 27.00 | 35.02 | 24.14 | 28.46 | 31.40 | 36.27 | 47.28 |
| | 100 | 17.14 | 17.14 | 23.35 | 27.04 | 35.02 | 21.14 | 28.46 | 31.40 | 36.33 | 47.28 |
| | 1000 | 17.14 | 17.14 | 23.41 | 27.05 | 35.02 | 21.14 | 28.45 | 31.40 | 36.31 | 47.28 |

*^♦ were taken from the output of GNU time. *User time was taken from “User time (seconds)” and converted to the nearest minute. ^Elapsed time was taken from “Elapsed (wall clock) time (h:mm:ss or m:ss)”. ♦Memory usage was taken from “Maximum resident set size (kbytes)” and converted to GB. ♥Only 1 thread was used. All other values shown are for 12 threads.
